# Effects of NAD⁺ Repletion with Nicotinamide Riboside on Obesity-Induced Chronic Kidney Disease and Renal Cell Lipotoxicity

**DOI:** 10.1101/2025.04.22.649334

**Authors:** Morgane Decarnoncle, Louise Pierre, Pauline Rouzé, Inès Jadot, Anne-Emilie Declèves, Florian Juszczak

## Abstract

Nicotinamide riboside (NR), a natural precursor of NAD⁺, has been suggested to confer protection against metabolic and age-related disorders. However, its impact on chronic kidney disease (CKD), particularly in the context of obesity, remains poorly understood. Here, we evaluated the potential effects of NR supplementation in models of obesity-induced renal injury. The metabolic and renal effects of both preventive and interventional NR supplementation were assessed in mice fed high-fat or low-fat diets. Our data showed that NAD⁺ repletion, whether preventive or interventional, did not affect body or organ weights, glucose metabolism, insulin resistance, or hepatic and renal lipid accumulation. NR supplementation was however associated with SIRT3-mediated deacetylation of SOD2 in the renal tissue of obese mice, and it moderately reduced renal dysfunction. To further explore the cellular mechanisms underlying the renal effects of NR in a lipotoxic context, we investigated its impact on renal proximal tubular cells exposed to palmitic acid (PA). NR significantly prevented oxidative stress in proximal tubular epithelial cells, as evidenced by the activation of SOD2 and the reduction of lipid peroxidation and mitochondrial dysfunction. However, NR did not reduce PA-induced lipid accumulation. In conclusion, this study provides evidence that NR exerts antioxidant effects and enhances mitochondrial function in renal cells in vitro but does not protect obese mice from metabolic disorders and associated CKD.

## Introduction

In last decades, obesity has escalated into a global epidemic, driven primarily by sedentary lifestyles and high-calorie diets.^1^ Obesity is a well-established risk factor for several comorbidities including abdominal adiposity, hyperlipidemia, hyperglycemia, hyperinsulinemia, and hypertension, which collectively contribute to the development of chronic diseases.^2^ Particularly, obesity is a significant risk factor for chronic kidney disease (CKD) and a strong predictor for the progression to end-stage renal disease (ESRD).^3^ Obesity-induced CKD is characterized by specific pathological changes including obesity-related glomerulopathy, defined by glomeruli enlargement, hypertension and hyperfiltration which may lead to the development of focal segmental glomerulosclerosis.^4^ These structural changes impair kidney function over time. Additionally, in rodents and humans, obesity has been shown to promote ectopic lipid accumulation in renal cells and particularly in proximal tubular epithelial cells (PTECs).^5–9^ This lipid accumulation disrupts normal lipid metabolism by inhibiting fatty acid oxidation (FAO) and activating lipogenesis. Overall, abnormal lipid metabolism in the kidneys has been shown to impair cell function and contribute to renal damage and CKD development.^5–9^ Moreover, evidence suggests that obesity disrupts mitochondrial function and increases oxidative stress in PTECs, contributing to renal damage.^10^ Particularly, these cells primarily rely on free fatty acids as an energy source, which are catabolized through mitochondrial FAO and oxidative phosphorylation (OXPHOS). Impaired mitochondrial function in PTECs leads to increased production of reactive oxygen species (ROS), which damage cellular components and promote inflammation.^11^

Despite extensive efforts to develop therapeutic strategies for obesity-related CKD, effective treatments remain limited, often accompanied by significant side effects. This highlights the urgent need for alternative therapies. Among them, those protecting kidney cells by supporting mitochondrial function and reducing oxidative stress seem to offer the potential to slow down the progression of kidney disease in obese individuals.^12^ Recently, studies have demonstrated that nicotinamide adenine dinucleotide (NAD^+^) metabolism is disturbed in human kidneys as well as in animal models with CKD.^13–15^ NAD⁺ is a coenzyme that plays a crucial role in mitochondrial ATP production and oxidative reactions. It also serves as a co-substrate for NAD⁺-dependent deacetylases, such as sirtuins. Sirtuins regulate key metabolic processes involved in maintaining cellular redox balance and mitochondrial integrity, including anti-oxidative response, mitochondrial biogenesis, mitochondrial dynamics and mitophagy.^16^ Particularly, sirtuin 3 (SIRT3) deacetylates and activates superoxide dismutase 2 (SOD2), a critical enzyme that mitigates oxidative damage by converting superoxide radicals into hydrogen peroxide and molecular oxygen.^17^ SIRT3 enhances the activity of SOD2 through deacetylation of specific lysine residues (such as K68).^17–19^ Mice deficient in quinolinate phosphoribosyltransferase (QPRT), an enzyme essential for *de novo* NAD^+^ biosynthesis, showed increased susceptibility to acute kidney injury (AKI).^14^ During AKI, both NAD^+^ consumption and synthesis are disrupted, leading to impaired FAO, ectopic lipid accumulation within the kidney, and tubular dysfunction.^20^ Additionally, diabetic kidney disease is linked to a reduction in the intrarenal NAD^+^/NADH ratio, which contributes to mitochondrial oxidative stress.^21^ These findings highlight the critical role of NAD⁺ in maintaining kidney function, particularly under conditions of stress such as acute kidney injury and diabetic kidney disease. While NAD^+^ depletion is linked to a wide spectrum of pathological conditions including neurological and age-related disorders, studies indicate that restoring NAD^+^ levels with precursors may alleviate these pathological conditions, potentially by supporting redox balance and mitochondrial function during cellular stress and injury.^22^ Nicotinamide Riboside (NR) is a form of vitamin B3 and a naturally occurring precursor of NAD^+^, found in sources such as cow’s milk.^23^ NR is converted to NAD⁺ primarily via the NR kinase (NRK) pathway.^24,25^ In mice and humans, NR treatment has been shown to be more efficient in boosting NAD^+^ than other NAD^+^ precursors such as Nicotinic Acid (NA) or Nicotinamide (NAM).^26^ Several studies have shown that NR supplementation increases NAD^+^ level, enhances mitochondrial biogenesis and oxidative metabolism, and thus protects against neurodegenerative disorders and age-related physiological decline in mammals.^27^ In their study, Canto and colleagues demonstrated that NR treatment at a dose of 400 mg/kg/day for 10 weeks in high-fat diet (HFD)-fed mice increased plasma and intracellular NAD⁺ levels in muscles, brown adipose tissue, and liver. NR supplementation also enhanced SIRT3-mediated SOD2 deacetylation in cultured cells and in the muscle tissue, which improved mitochondrial function and oxidative capacity.^28^ However, the particular effects of NR supplementation for the renal tissue and obesity-associated kidney dysfunction is still unknown. This study investigated the effects of NR treatment in obese mice with CKD as well as in PTECs in conditions of lipotoxicity. We showed that, *in vitro*, NR supplementation reduces oxidative stress and enhances mitochondrial function in PTECs. However*, in vivo*, it does not prevent renal dysfunction and instead induces lipid accumulation in both the kidney and liver, particularly with long-term supplementation.

## Results

### Preventive or interventional NR supplementation does not impact metabolic syndrome parameters

We investigated the effect of early and late NR supplementation in low-fat diet (LFD) or high-fat diet (HFD) mice. NR was administered either concomitantly with the diet (LFD NR20 and HFD NR20) or as an interventional treatment during the last 8 weeks of the protocol (LFD NR8 and HFD NR8) (**Figure 1A**). Indeed, obese mice fed a HFD for at least 12 weeks are known to develop characteristic features of obesity-related nephropathy.^29^ The data presented in **Figure 1B** indicate that when comparing mice fed a HFD to those fed a LFD, there was a significant increase in body weight starting from week 8. However, when comparing mice within the same dietary groups, no significant differences in body weight were observed as a result of NR supplementation. Moreover, the weights of kidneys and the liver were significantly higher in all HFD-fed mice groups compared with LFD-fed mice groups (**Figure 1C**, **D**), while the heart weight was similar (**Figure 1E**). In accordance with previous reports, mice fed a HFD for 20 weeks developed glucose intolerance, hyperglycemia, and hyperinsulinemia.^8^ The glycemia and insulinemia were significantly higher in all HFD groups compared to LFD groups with no significant effect of preventive or interventional NR supplementation (**Figure 2D**, **E**). Moreover, results of the glucose tolerance test (GTT) at week 0, 12 and 20 confirmed the development of glucose intolerance in HFD animals and the absence of NR supplementation effect (**Figure A-C**). The calculation of the HOMA-IR (Homeostatic Model Assessment for Insulin Resistance), an indicator of insulin resistance, demonstrated that NR supplementation had no effect on IR in HFD mice (**Figure 2F**). Altogether, these results show that NR has no influence on impaired glucose metabolism and insulin resistance in obese mice. The plasma levels of cholesterol were also measured in each experimental group as another indicator of the metabolic syndrome (**Figure 2G**). As illustrated, HFD induced a significant elevation of plasma cholesterol levels, whereas this increase was counteracted only in the HFD NR20 group compared to HFD. Furthermore, we investigated plasma interleukin 6 (IL-6) as a marker of systemic inflammation and the data reported an increased concentration of this proinflammatory cytokine in HFD mice that was not significantly reduced with NR supplementation (**Figure 2H**).

**Figure 1.**
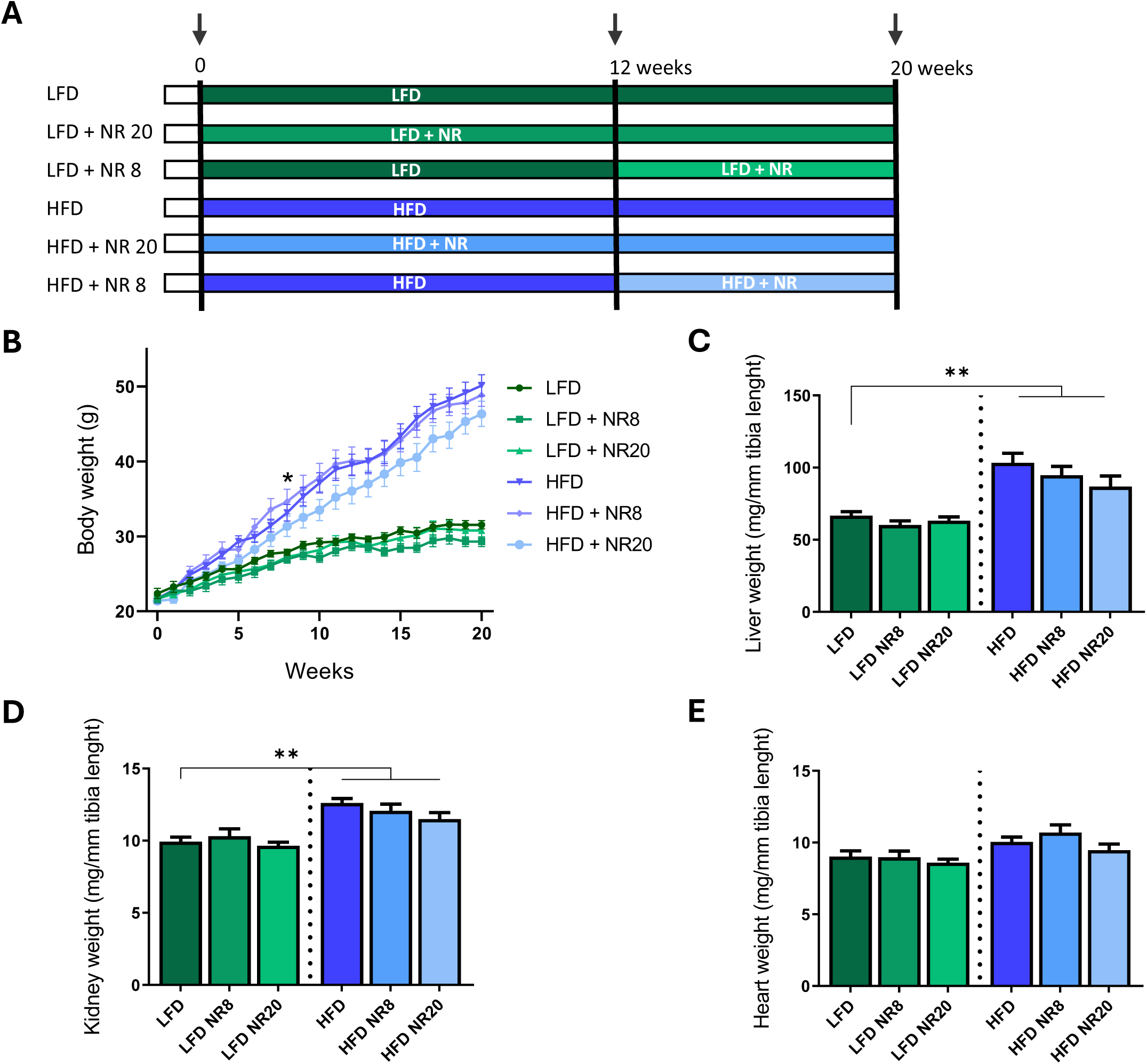
Preventive or interventional NR supplementation did not impact body nor organ weights in mice fed with low- or high-fat diet. (**A)** Schematic representation of experimental design. Eight-weeks old C57Bl6/J male mice were randomized to either a low- (LFD) or high-fat diet (HFD) supplemented or not with 400 mg/kg/Day of nicotinamide riboside (NR) from week 0 (preventive) or week 12 (interventional) for a total of 20 weeks. At week 0, 12, and 20 (represented by black arrows), urine collection and glucose tolerance test were performed. **(B)** Body weight evolution over time of LFD or HFD animals supplemented or not with NR. **(C-E)** Measurements of the **(C)** liver, **(D)** kidney, and **(E)** heart weights at week 20 in LFD or HFD animals supplemented or not with NR. Data are presented as means ± SEM, n=10 in each group. Statistical analyses were performed by two-way ANOVA followed by Sidak’s post hoc test. *p ≤ 0.05; **p ≤ 0.01 between groups.

**Figure 2.**
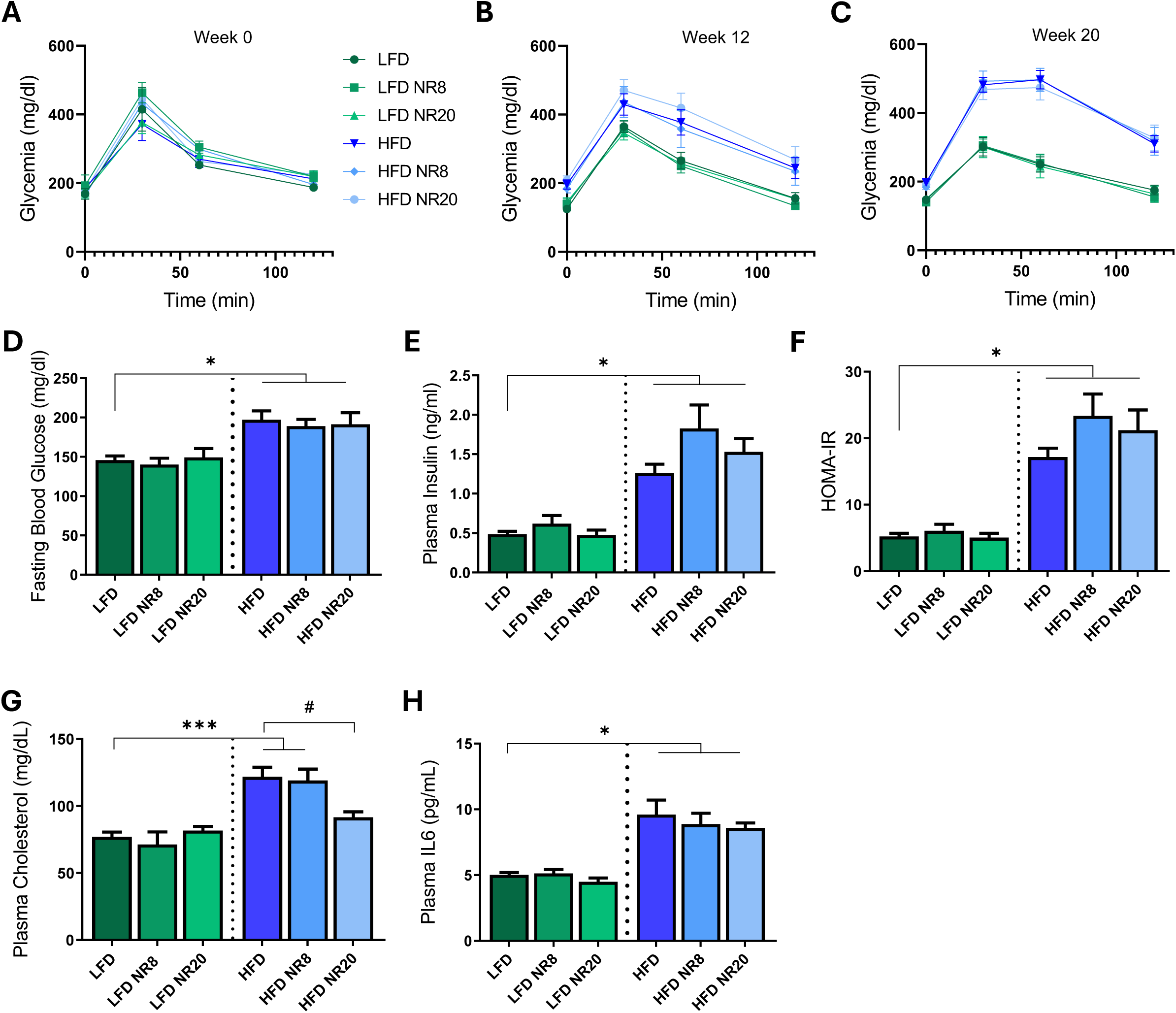
Preventive or interventional NR supplementation did not improve parameters of the metabolic syndrome. **(A-C)** Glucose tolerance tests of LFD or HFD mice supplemented or not with NR at **(A)** week 0, **(B)** week 12 and **(C)** week 20. Glycemia was measured in fasted mice before (0 min) and 30, 60 and 120 min after intraperitoneal injection of 2 g/kg b.w. glucose. **(D)** Measurement of fasted blood glucose at week 20 in LFD or HFD animals supplemented or not with NR. **(E)** Measurement of plasma insulin at week 20 in LFD or HFD animals supplemented or not with NR. **(F)** HOMA-IR index for LFD or HFD animals supplemented or not with NR after 20 weeks. **(G)** Assessment of plasma cholesterol levels at week 20 in LFD or HFD animals supplemented or not with NR. **(H)** Measurement of plasma IL6 levels at week 20 in LFD or HFD animals supplemented or not with NR. Data are presented as means ± SEM, n=10 in each group. Statistical analyses were performed by two-way ANOVA followed by Sidak’s post hoc test. *p ≤ 0.05; ***p ≤ 0.001; #p ≤ 0.05 between groups.

### NR supplementation does not improve hepatic steatosis in obese mice but reduces liver fibrosis

HFD-fed mice exhibited features of hepatic steatosis, characterized by substantial lipid droplet (LD) accumulation within hepatocytes (**Figure 3A**, **B**). Notably, NR supplementation did not alleviate hepatic lipid accumulation in HFD-fed mice. Interestingly, moderate LD accumulation was observed in hepatocytes of LFD animals receiving NR, regardless of the supplementation duration. In addition, hepatic fibrosis was assessed using Sirius Red staining. As shown in **Figure 3C**, liver tissues from HFD-fed mice displayed a significantly higher percentage of fibrosis compared to the LFD control groups. However, both early and late NR supplementation effectively prevented the development of liver fibrosis in HFD-fed mice. Thus, while HFD-fed mice exhibited marked hepatic fibrosis, NR supplementation prevented fibrosis development, highlighting its potential role in mitigating diet-induced liver damage.

**Figure 3.**
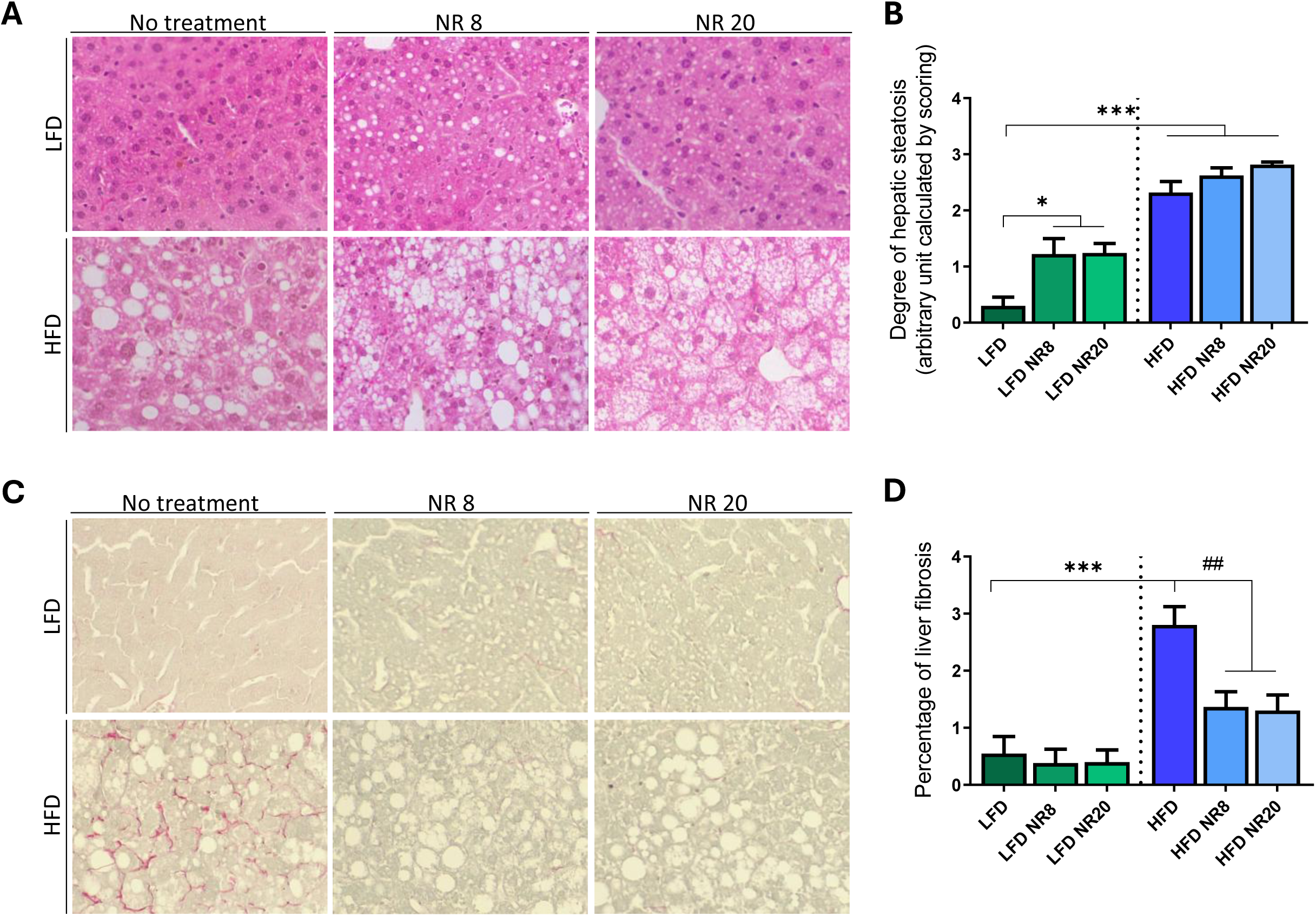
NR supplementation reduces liver fibrosis but does not improve hepatic steatosis in obese mice. (**A)** Representative micrographs of liver sections from LFD or HFD animals supplemented or not with NR at week 20 and stained with hematoxylin and eosin (original magnification ×400). **(B)** Semi-quantitative quantification of hepatic steatosis by scoring. **(C)** Representative micrographs liver sections from LFD or HFD animals supplemented or not with NR at week 20 and stained with Sirius Red (original magnification ×400). **(D)** Quantitative analysis of the percentages of staining positive for Sirius red indicative of liver fibrosis. Data are presented as means ± SEM, n=10 in each group. Statistical analyses were performed by two-way ANOVA followed by Tukey’s post hoc test. *p ≤ 0.05; ***p ≤ 0.001; ##p ≤ 0.01 between groups.

### NR supplementation does not reduce renal lipid accumulation and is associated with moderate improvement of renal function

HFD mice typically exhibit significant alterations in renal function and disturbance in lipid metabolism.^7^ HFD is associated with glomerular hypertrophy, mesangial matrix expansion, and a decline in renal function, as evidenced by increased albuminuria.^8^ Additionally, LD accumulation is commonly observed in proximal tubular cells of HFD-fed animals, reflecting lipid metabolism dysregulation in the kidney.^5,7^ **Figure 4** demonstrates that mice fed a HFD alone developed glomerular hypertrophy as evidenced by a slight increase (p=0.0594) in mesangial matrix expansion (**Figure 4A-C**). It is concomitant with renal function decline in HFD animals as they present a significantly increased albuminuria (**Figure 4D**). Interestingly, in HFD mice supplemented with NR as an interventional treatment, the albuminuria is decreased compared to HFD mice but not as a preventive strategy (**Figure 4D**). We further examined LDs accumulation in proximal tubules. We confirmed the presence of LD in PTECs in HFD mice, while NR supplementation had no effect on this parameter (**Figure E, F**). Similarly to liver tissue, LFD group treated with NR for 20 weeks present a moderate but significantly increased LD accumulation in proximal tubules. Canto and colleagues demonstrated that NR supplementation induces SIRT3-dependant deacetylation of SOD2 in muscle tissue of obese mice.^28^ To address whether SIRT3 was effectively activated in renal tissue by NR supplementation in LFD- and HFD-fed mice, we analyzed the abundance of acetylated SOD2 and lysine-acetylated proteins in the mitochondrial-enriched fraction of the protein levels, but this did not reach statistical significance (**Figure 4G-H**). However, a significant increase in acetylated-SOD2 abundance was observed in HFD-fed animals compared to LFD-fed ones (**Figure 4G**, **I**). As expected, it suggests a decreased SIRT3 activity in mitochondria. Moreover, preventive and interventional NR treatment in HFD-fed mice significantly reduced the acetylation level of SOD2, confirming SIRT3 activation in response to NR in the renal tissue of obese mice.

**Figure 4.**
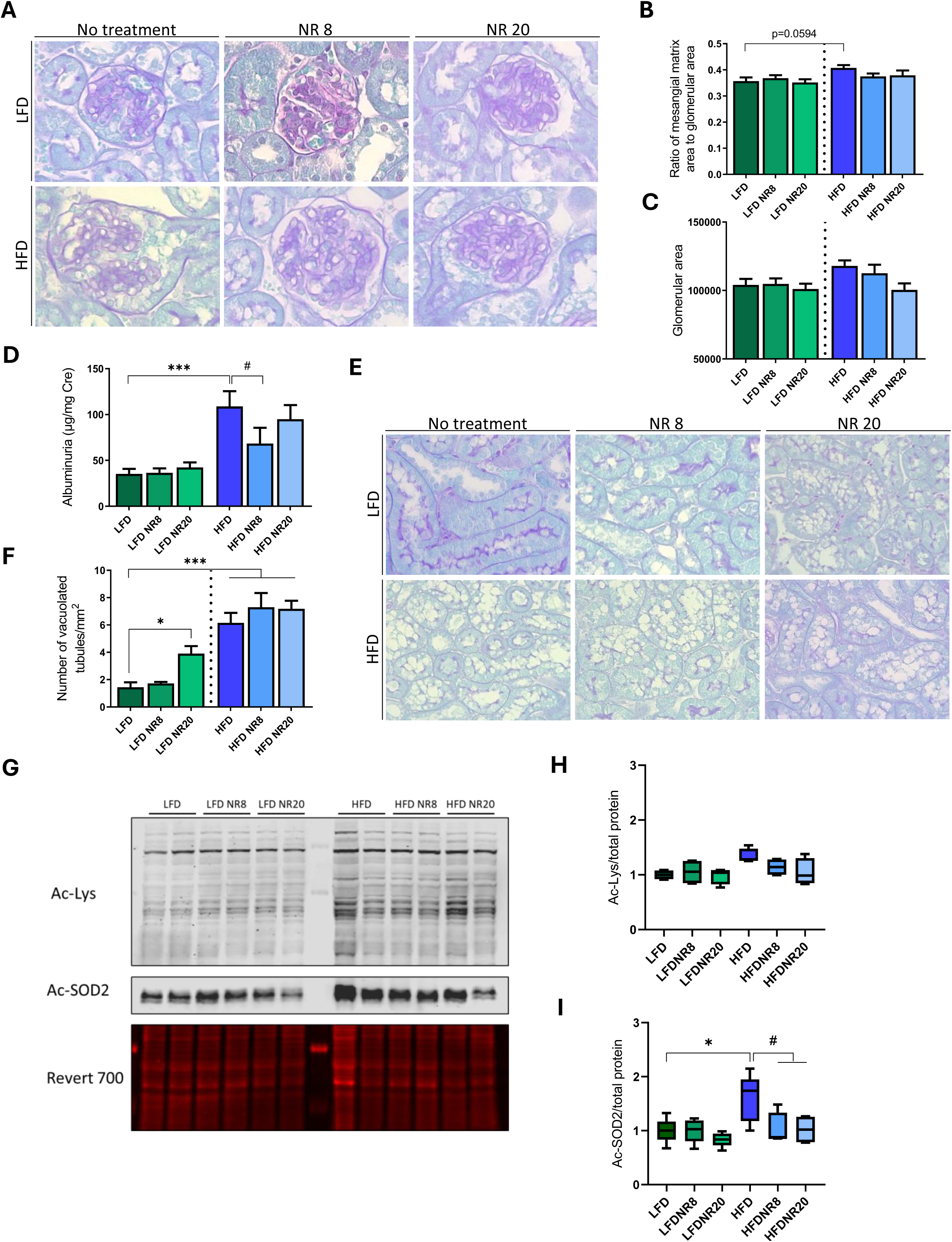
NR supplementation has limited effects on renal function and does not reduce ectopic lipid accumulation in proximal tubules of obese mice. **(A)** Representative micrographs of kidneys sections from LFD or HFD animals supplemented or not with NR at week 20 and stained with of Periodic Acid Schiff and Luxol Fast Blue (original magnification ×400). **(B, C)** Quantification of the ratio of mesangial matrix area to glomerular area and the glomerular area from 15 glomeruli per kidney section with one section per animal. **(D)** Quantitative measurement of the ratio between urinary albumin and urinary creatinine at week 20 in LFD or HFD animals supplemented or not with NR at week 20. **(E)** Representative micrographs of kidneys sections from LFD or HFD animals supplemented or not with NR at week 20 and stained with Periodic Acid Schiff and Luxol Fast Blue (original magnification ×400). **(F)** Quantitative analysis of number of vacuolated tubules per mm^2^ of renal section. **(G)** Data are presented as means ± SEM, n=10 in each group. Representative western blot and **(H, I)** quantitative densitometry analysis of acetylated lysine (Ac-Lys), acetylated SOD2 (Ac-SOD2, K68 residue) and total protein content (Revert 700) in mitochondria-enriched fractions from renal cortex of LFD or HFD animals supplemented or not with NR at week 20. Data is presented as means ± SEM, n=4-6 in each group. Statistical analyses were performed by two-way ANOVA followed by Sidak’s post hoc test. *p ≤ 0.05; ***p ≤ 0.001; #p ≤ 0.05 between groups.

### NR supplementation protects human PTECs from PA-induced metabolic changes but not from LD accumulation

Given that NR supplementation did not prevent renal dysfunction *in vivo* but increased lipid accumulation, we aimed to further investigate its effects at the cellular level. Specifically, we aimed to better characterize palmitic acid (PA)-induced lipotoxicity in PTECs and determine whether NR could still exert protective effects by restoring NAD⁺ levels. Using an *in vitro* model, we assessed whether NR could counteract PA-induced metabolic dysfunctions in human PTECs, independently of systemic factors. Different PA concentrations have been applied on HK-2 cells (human PTECs line) to mimic metabolic dysfunctions observed in obese mice. Therefore, HK-2 cells were treated from 100 to 700 µM of PA for 24h (**Supp. Figure 1**). As observed, 300 µM of PA for 24h led to a decreased NAD^+^/NADH ratio and is associated with an increased positive staining for Oil Red O and BODIPY in HK-2, suggesting an accumulation of neutral lipids. Thus, regarding our data and previous studies, we decided to use a 300µM concentration of PA for the following experiments. We next confirmed that a significant increase in NAD^+^/NADH ratio was observed with 1 mM NR (**Figure 5A**). Then, we would like to determine whether NR-mediated restoration of the NAD^+^/NADH ratio could protect HK-2 cells from deleterious consequences of PA-induced lipotoxicity. To address this question, HK-2 cells were exposed to 0.4% BSA as a vehicle or 300 µM PA with or without 1 mM NR for 24 h. NR significantly increases the NAD^+^/NADH ratio in BSA- and PA-treated cells (**Figure 5B**). NR treatment did not significantly affect the number nor size of LD in PA-treated HK-2 cells (**Figure 5C-E**), even if it significantly increased their metabolic activities (**Figure 5F**).

**Figure 5.**
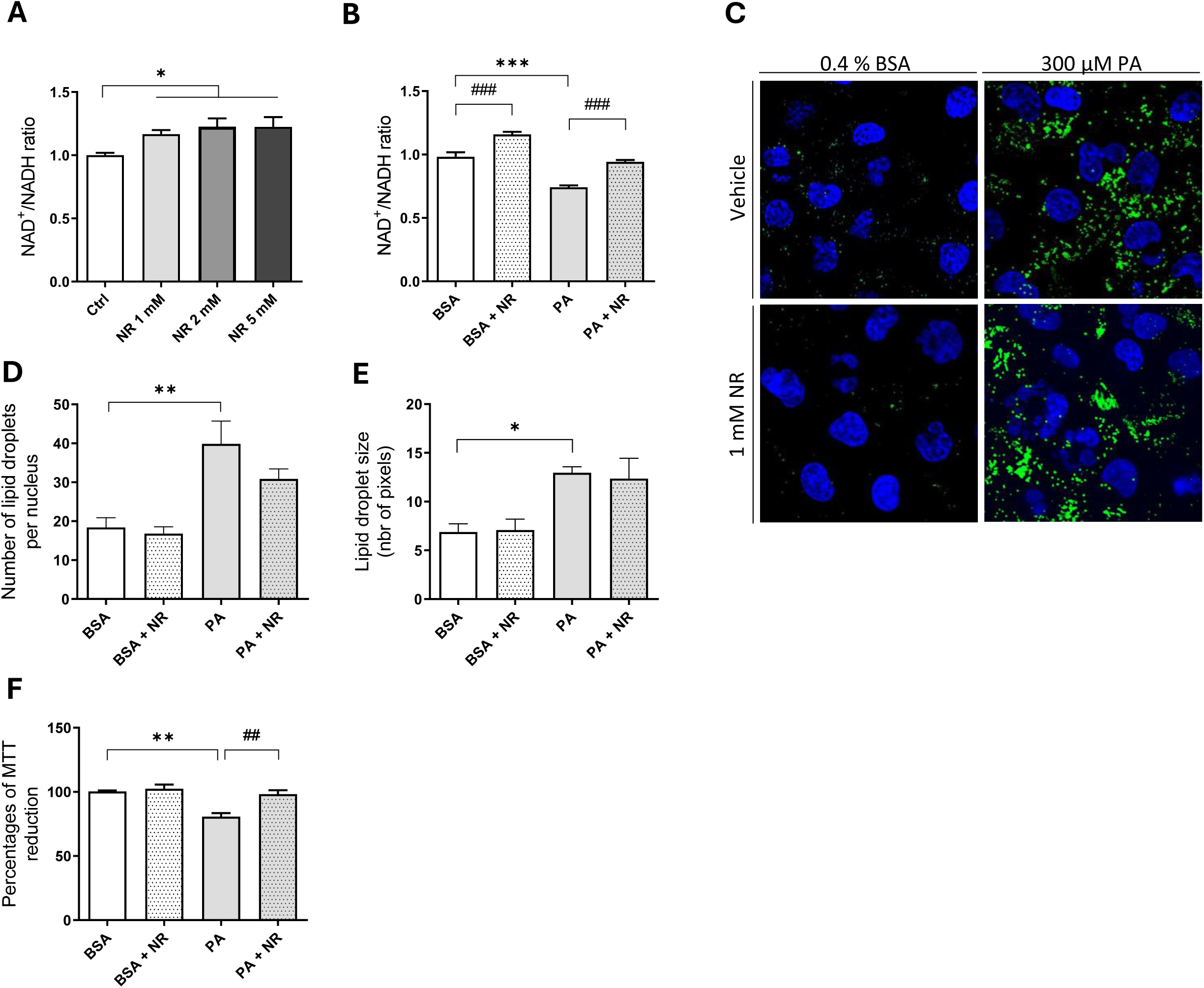
NR treatment protects HK2 cells from PA-induced metabolic changes but does not prevent lipid droplet formation. **(A)** Measurement of the NAD^+^/NADH ratio using the NAD/NADH-Glo Assay Kit in HK2 cells treated for 24 h with 1-, 2- or 5-mM NR or vehicle. **(B)** Measurement of the NAD^+^/NADH ratio using the NAD/NADH-Glo Assay Kit in HK2 cells treated for 24 h with 0.4 % BSA or 300 µm PA with or without 1 mM NR. (C) Representative fluorescent micrographs of HK2 cells treated for 24 h with 300 µM PA or 0.4 % BSA with or without 1 mM NR and stained with 2.5 µM BODIPY^TM^ 493/503 for 15 min (400X). **(D, E)** Quantifications of lipid droplet number and size on 100 cells per condition by Lipid Droplets MRI tool. **(F)** Percentages of MTT reduction (indicative of metabolic activities) in HK2 cells treated for 24 h with 300 µM PA or 0.4 % BSA with or without 1 mM NR. Data are presented as means ± SEM of three independent biological experiments. Statistical analyses were performed by **(A)** one-way ANOVA followed by Dunnett’s post hoc test or by **(B, D, E, F)** two-way ANOVA followed Tukey’s post hoc test. *p ≤ 0.05 ; **p ≤ 0.01; ***p ≤ 0.001; ###p ≤ 0.01; ###p ≤ 0.001 between groups.

### NR reduces PA-induced mitochondrial oxidative stress and lipid peroxidation in PTECs

ROS are short-lived oxygen-containing molecules known for their high reactivity, often promoting oxidative stress. These molecules are natural byproducts of aerobic respiration and primarily originate from mitochondria. PTECs are highly dependent on mitochondrial functions to sustain their metabolism, making them particularly vulnerable to mitochondrial damage.^30^ PA, a saturated fatty acid, is well-documented for inducing oxidative stress, further exacerbating mitochondrial dysfunction in these metabolically demanding cells. As expected, PA is associated with a significantly increased MitoSOX fluorescence indicative of mitochondrial ROS compared to BSA in HK-2 cells (**Figure 6A**, **B**). However, this increased mitochondrial superoxide production was prevented upon NR supplementation in PA-treated cells. To confirm the benefits of NR on oxidative balance, its effect on lipid peroxidation was then addressed with a lipid peroxidation sensor that shifts from red to green in presence of lipid hydroperoxides. The green-to-red fluorescence ratio was significantly increased in response to PA, indicating enhanced lipid peroxidation **(Figure 6C, D)**. This effect was reversed by NR treatment. We further evaluated the abundance of acetylated-SOD2 under these different experimental conditions (**Figure 6E**, **F**). Consistent with our *in vivo* findings, acetylated-SOD2 levels are higher in the PA condition compared to the control group. However, a significant decrease in acetylated-SOD2 is observed in cells treated with both PA and NR, compared to those treated with PA alone. This confirms that NR induces SIRT3-mediated antioxidative response in renal cells.

**Figure 6.**
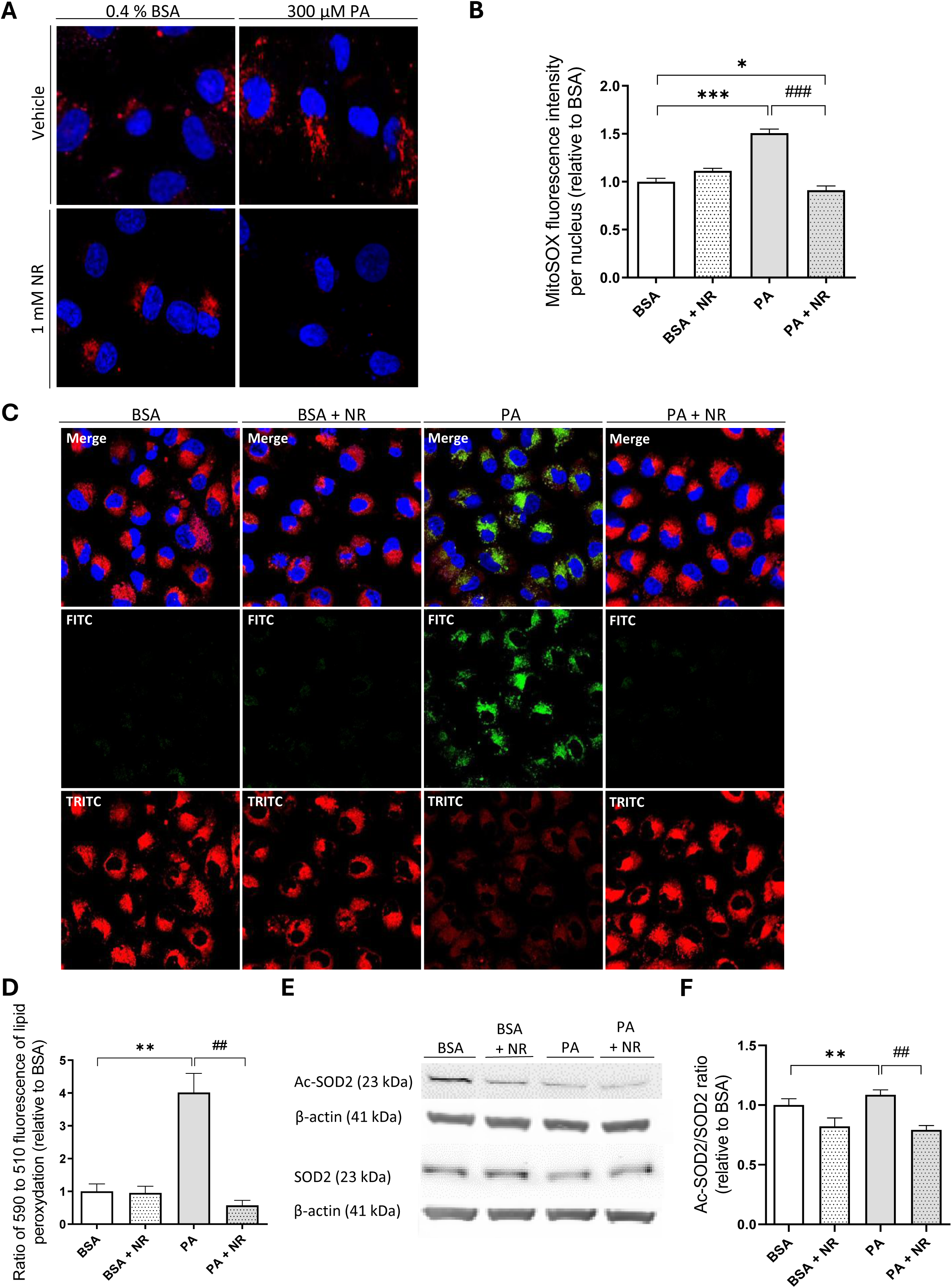
NR treatment prevents PA-induced oxidative stress and lipid peroxidation in HK2 cells. **(A)** Representative fluorescent micrographs of HK2 cells treated for 24 h with 300 µM PA or 0.4 % BSA with or without 1 mM NR and stained with 5 µM MitoSOX^TM^ for 15 min (400X). **(B)** Quantification of the MitoSOX-relative fluorescence intensity per nucleus on more than 100 cells per experimental condition. **(C)** Representative fluorescent micrographs of HK2 cells treated for 24 h with 300 µM PA or 0.4 % BSA with or without 1 mM NR and stained with 10 nM Image-iT^TM^ Lipid Peroxidation for 15 min (400X). **(D)** Quantitative analysis of the 590 to 510 nm ratio of fluorescence intensities linked to Image-iT^TM^ Lipid Peroxidation staining on more than 100 cells per experimental condition. **(G)** Representative western blot and **(H)** quantitative densitometry analysis of acetylated SOD2 (Ac-SOD2, K68 residue), SOD2 and β-actin in in HK2 cells treated for 24 h with 300 µM PA or 0.4 % BSA with or without 1 mM NR. Data are presented as means ± SEM of three independent biological experiments. Statistical analyses were performed by two-way ANOVA followed Tukey’s post hoc test. *p ≤ 0.05 ; **p ≤ 0.01; ***p ≤ 0.001; ##p ≤ 0.01; ###p ≤ 0.001 between groups.

### NR prevents PA-induced mitochondrial damage in PTECs

Given that NR induces an antioxidant response mediated by SOD2, we hypothesized that it could impact the integrity of the mitochondrial network, protect mitochondrial activity, and support mitochondrial dynamics. We thus assessed the potential benefits of NR on the morphology of the mitochondrial network in PA-treated cells. MitoTracker Green, which accumulates in mitochondria independently of their potential, was used to stain the mitochondrial network and to calculate the form factor, the ratio between endpoints to branched indicating network interconnectivity (**Figure 7A**, **B**). BSA-treated cells displayed a filamentous and interconnected network while it appeared fragmented in PA-treated cells. This event was confirmed by a significant increased form factor (mitochondrial endpoints to branched) in response to PA. Notably, PA-induced mitochondrial fragmentation was prevented by NR as the form factor was significantly reduced in cells treated with PA combined with NR compared to PA alone. We next asked whether NR may be beneficial against PA-induced mitochondrial fragmentation by preventing an underlying event, mitochondrial depolarisation. HK-2 cells treated with BSA or PA with or without NR were thus stained with both the MitoTracker Red dye, which stains mitochondrial with normal value of potential as well as the ratiometric JC-1 dye (**Figure 7C-D**). PTECs treated with PA displayed a decreased both MitoTracker Red depolarisation which was prevented by NR. Mitochondrial DNA (mtDNA) to nuclear DNA (nDNA) ratio was used to evaluate mitochondrial abundance. Surprisingly, mitochondrial content (mtDNA)/nDNA) was decreased with NR supplementation, but the RNA expression of the Mitochondrial transcription factor A (*TFAM*) and BCL2 Interacting Protein 3-like (*BNIP3L*) increased in PA condition treated with NR (**Figure 7E-G**). Elevated *BNIP3L* expression indicates enhanced mitophagy, a process where damaged or dysfunctional mitochondria are selectively degraded. This could explain the reduction in mtDNA/nDNA, as NR might promote the clearance of impaired mitochondria. Increased *TFAM* expression suggests that NR is promoting mitochondrial biogenesis. However, biogenesis may not yet fully compensate for the mitophagy-induced reduction in total mtDNA content, reflecting a dynamic mitochondrial turnover. Thus, our data suggests that NR improves mitochondrial quality by balancing mitophagy and biogenesis, thereby promoting the replacement of damaged mitochondria.

**Figure 7.**
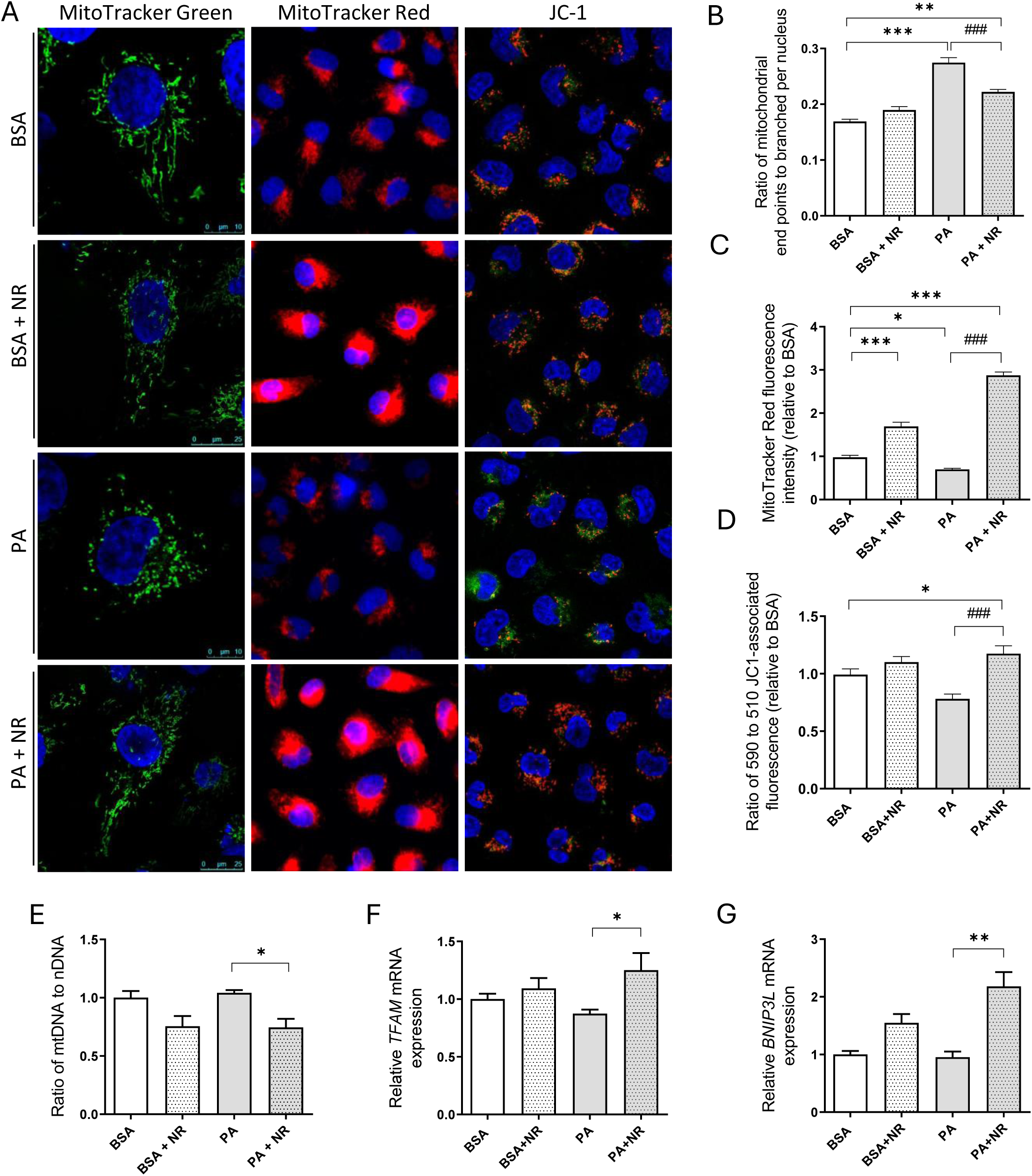
NR treatment prevented PA-induced mitochondrial damage in HK2 cells. **(A)** Representative fluorescent micrographs (400X) of MitoTracker™ Green FM, MitoTracker^TM^ Red CMXRos and MitoProbe™ JC-1 probe staining in HK2 treated with BSA 0,4%, PA 300 µM with or without NR 1 mM. Nuclei were counterstained with Hoechst Dye (blue). **(B)** Quantification of ratio between mitochondrial end points to branched on more than 100 cells per experimental condition. **(C)** Quantitative analysis of MitoTracker^TM^ Red CMXRos fluorescence intensities on more than 100 cells per experimental condition. **(D)** Quantitative analysis of the 590 to 510 nm ratio of JC-1 fluorescence intensities on more than 100 cells per experimental condition. **(E)** Quantitative analysis of the mitochondrial (mt) to nuclear (n) DNA ratio from HK2 cells treated for 24 h with 300 µM PA or 0.4 % BSA with or without 1 mM NR. **(F,G)** Relative mRNA expression of *TFAM* and *BNIP3L* in HK2 cells treated for 24 h with 300 µM PA or 0.4 % BSA with or without 1 mM NR. Data are presented as means ± SEM of three independent biological experiments. Statistical analyses were performed by two-way ANOVA followed Tukey’s post hoc test. *p ≤ 0.05 ; **p ≤ 0.01; ***p ≤ 0.001; ##p ≤ 0.01; ###p ≤ 0.001 between groups.

## Discussion

Several studies have investigated the beneficial effects of NAD^+^ precursors in mitigating diet-induced obesity.^28,31,32^ However, to the best of our knowledge, no investigation have specifically explored the impact of NR supplementation on the kidneys in conditions of obesity and lipotoxicity. Both decreased NAD^+^ biosynthesis pathway and NAD^+^ consumption are responsible for NAD^+^ reduction in AKI and diabetic kidney disease (DKD).^33,34^ In AKI, endoplasmic reticulum stress response reduces *de novo* NAD^+^ biosynthesis in the kidney by reducing QPRT expression.^35^ Persistent QPRT reduction, a key enzyme of *de novo* NAD^+^ biosynthesis, is associated with AKI to CKD progression. In DKD, reduced NAD^+^ levels have been linked to decreased activity of the enzyme kynurenine 3-monooxygenase (KMO), also involved in the NAD^+^ biosynthesis pathway.^34^ Additionally, NAD^+^-dependent enzymes, such as sirtuins and poly(ADP-ribose) polymerases (PARPs) consume NAD^+^ during DNA repair and stress responses, further contributing to NAD^+^ depletion in both AKI and diabetes.^16^ NAD^+^ precursors such as Nicotinamide Mononucleotide (NMN) and NAM have been reported beneficial effects in renal injury by restoring NAD^+^ levels, enhancing mitochondrial function, reducing oxidative stress and inflammation, preventing cellular senescence and fibrosis, and improving tubular cell function and survival.^36–39^ In this study, we aimed to evaluate the effects of NR-mediated NAD^+^ repletion in HFD-fed mouse model of CKD, previously established and characterized by our group.^8,9,40^ NR has been found to be readily absorbed, to enhance NAD^+^ levels more effectively than other precursors, and to exhibit superior bioavailability as a form of vitamin B3 in both mice and humans.^26^ Here, NR was used either as a preventive therapy (early administration of NR in the diet for 20 weeks) or as a treatment (delayed administration in the diet for the last 8 weeks of the protocol). Canto and colleagues demonstrated that 400 mg/kg/Day of NR treatment in HFD mice for 10 weeks increased plasma and intracellular NAD^+^.^41^ Here, we demonstrated that 400 mg/kg/Day of NR supplementation in the diet neither early or late NR were associated with body weight loss or improvement of glucose intolerance, insulin resistance and inflammation. Several other studies have reported limited or no effect of NR supplementation on insulin sensitivity. Cartwright and colleagues demonstrated that while 8 weeks NR supplementation (500 mg/kg/day in drinking water) increased mitochondrial respiration in the muscle tissue, it did not counteract HFD-induced metabolic parameters, including glucose intolerance and hepatic lipid accumulation.^42^ Moreover, in another study, NR supplementation (350 mg/kg/Day in the diet) for 14 weeks in mice fed a HFD since week 15 did not altered body weight or glucose tolerance but moderately improved insulin resistance.^43^ Here, plasma cholesterol levels decreased following 20 weeks of NR supplementation in obese mice. However, this was not accompanied by a reduction in the liver steatosis score. More surprisingly, NR treatment in control mice was associated with an increased lipid deposition in the liver tissue. In contrast, NR was associated with beneficial effects regarding liver fibrosis as attested by collagen deposition that was markedly reduced in HFD mice supplemented with NR. Pham and colleagues reported the effects of NR supplementation for 20 weeks (400 mg/kg/Day in the diet) on hepatic fibrosis in mice fed a high-fat/high-sucrose/high-cholesterol diet.^44^ Interestingly, they showed that NR markedly reduced collagen accumulation in the liver but not lipid accumulation as we also observed in our study, suggesting that the protective effect of NR on liver fibrosis was independent of changes in liver steatosis. Next, we demonstrated for the very first time the ability of NR to activate SIRT3 in mitochondrial fraction of renal cortex as evidenced by the increased SIRT3-mediated deacetylation of SOD2. SIRT3 is a NAD⁺-dependent deacetylase primarily localized to mitochondria, where it regulates the activity of several metabolic and antioxidant enzymes through deacetylation. One of its well-established targets is SOD2, a mitochondrial enzyme critical for detoxifying ROS. The deacetylation of SOD2 by SIRT3 enhances its enzymatic activity, contributing to improved mitochondrial oxidative defense. Reduced SIRT3 activity and SOD2 hyperacetylation contribute to oxidative stress and associated mitochondrial dysfunction ^18^. HFD consumption has been shown to increase SOD2 acetylation.^45^ Canto and colleagues have demonstrated that NR induces SIRT3-mediated SOD2 deacetylation in the skeletal muscle, liver, and brown adipose tissue but not in brain and white adipose tissue.^41^ In our experimental model, HFD mice exhibited increased acetylation of mitochondrial SOD2 in renal tissue, that was completely reversed by NR supplementation. We also described the effects of NR on kidney function, glomerular and tubular injuries. The results indicate that NR supplementation in HFD mice has a limited impact on obesity-induced kidney injury. Specifically, NR induced only a moderate reduction in albuminuria, observed exclusively in the HFD group supplemented with NR for the last 8 weeks of the protocol. However, it did not significantly alter glomerular hypertrophy. Additionally, NR supplementation did not significantly reduce ectopic lipid accumulation in tubular cells, consistent with findings in the liver. Moreover, we further characterized the effects of NR on human PTECs (HK-2). Cells were exposed to PA to mimic obesity *in vitro*. PA is the most abundant saturated FA in the Western diet ^46^. Exposition with PA is widely used to investigate lipotoxicity on renal cell cultures.^47–49^ Here, NR treatment was associated with NAD⁺ repletion and an improvement in metabolic activity in PTECs exposed to PA. However, this was not accompanied by a reduction in intracellular LD, similarly to what is observed in the renal tissue of obese mice treated with NR. Moreover, the acetylation level of SOD2 was found to be decreased in response to NR, likely mediated by the activation of SIRT3. NR treatment was also associated with a reduction in mitochondrial ROS and the subsequent lipid peroxidation triggered by PA exposure. NR also counteracted the detrimental effects of PA on mitochondrial structure and integrity. This indicates that NR supports mitochondrial health and functionality, which are compromised under lipotoxic conditions. These improvements may underline the broader protective effects of NR against cellular oxidative stress and metabolic dysfunction in this context. Specifically, NR treatment alleviates the alterations in the mitochondrial network induced by PA. Moreover, NR reduces mitochondrial content in PTECs, not through decreased mitochondrial biogenesis but rather via increased mitochondrial degradation as evidenced by the upregulation of the mitophagy receptor BNIP3L. This observation is supported by a study showing that enhancing NAD^+^ levels with precursors promotes mitophagy, thereby maintaining mitochondrial homeostasis.^50^ In another study, NR supplementation was shown to elevate NAD⁺ levels, which in turn increased the expression of BNIP3L in human neurons derived from Parkinson’s disease patients.^51^ This upregulation of BNIP3L facilitated the clearance of damaged mitochondria through mitophagy, thereby ameliorating mitochondrial dysfunction associated with the disease. Lastly, a study found that increasing NAD^+^ levels via NR supplementation restored mitophagy in *C. elegans* model of accelerated aging through the worm homolog of BNIP3L.^52^

## Conclusion

Our study, in line with others, demonstrates that NR supplementation is well-tolerated but offers moderate beneficial effects on obesity-related conditions. However, its efficacy may vary depending on factors such as dose, treatment duration, animal source, administration method, and sex. Notably, emerging evidence indicates that high doses or prolonged NR treatment may cause adverse effects such as glucose intolerance and white adipose tissue dysfunction due to decreased metabolic flexibility in mice.^53,54^ Under our experimental conditions, NR yielded minimal improvements in metabolic parameters. It induced fat deposition in the kidney and liver in control animals, with more pronounced effects over extended treatment durations. Importantly, NR had no negative impact on kidney function and moderately ameliorated obesity-induced renal impairments, including reductions in albuminuria and oxidative stress. In proximal tubular cells, NR enhanced mitochondrial dynamics and triggered a robust antioxidant response, likely linked to SIRT3 activation, although it did not reduce lipid accumulation. Given ongoing clinical trials, further research is crucial to refine administration strategies and better understand potential adverse effects associated with NAD^+^ precursors.

## Material and methods

### Animals

The study conformed to the APS Guidelines for the Care and Use of Animals and was approved by the Animal Ethics Committee of the University of Namur. The experiments were conducted on C57Bl/6J male mice (Janvier Labs, Le Genest Saint-Isle, France) that were housed in cages with free access to food and water. Mice were maintained at 35–40 % relative humidity and a temperature of 20–23 °C with a 12:12 h light–dark cycle. Eight-weeks old mice were randomized to either a low-fat diet (LFD, 10% of total calories from fat; D12450J, Research Diets, New Brunswick, NJ, USA) or a high-fat diet (HFD, 60% of total calories from fat; D12492, Research Diets, New Brunswick, NJ, USA) supplemented or not with 400 mg/kg/Day of Nicotinamide Riboside (NR; Niagen, ChromaDex, Irvine, CA; prepared by Research Diets; D19052102 and D19052103) from week 0 or week 12 for a total of 20 weeks period. After 20 weeks, mice were euthanized, and blood samples were collected while kidneys were dissected into cortex and medulla. A kidney portion of cortex was freshly processed for mitochondrial isolation with the Mitochondria Isolation Kit for Tissue (Thermo Fisher Scientific, Waltham, MA, USA), following manufacturer’s instructions. Then, each renal sample was snap-frozen in liquid nitrogen for further analysis. An additional portion was fixed in 4% paraformaldehyde (PAF) for histological analysis. Mice were placed in metabolic cages for 24-h urine collection at baseline and at the end of the protocol. Urinary albumin and creatinine levels were measured using a mouse Albuwell ELISA kit and Creatinine Companion kit (Exocell, Philadelphia, PA, USA).

### Glucose Tolerance Test

After a 12 h-overnight fast, a glucose tolerance test (GTT) was performed at weeks 0, 12 and 20 of the experimental protocol. A dose of 2 g/kg body weight of D-glucose (Roth, Karlsruhe, Germany) was administered intraperitoneally. Blood samples were then obtained from the caudal vein, and the blood glucose level was measured at 0, 30, 60, and 120 min after glucose injection using a One Touch^®^ Verio glucometer (Zug, Switzerland).

### Biochemical assays

Plasma insulin levels were determined by ELISA using the rat/mouse insulin ELISA kit (Merck, Darmstadt, Germany). The homeostasis model assessment (HOMA-IR) for the insulin resistance index was determined using a calculator available from the Oxford Center for Diabetes, Endocrinology, and Metabolism (https://www.dtu.ox.ac.uk/homacalculator/). Plasma IL-6 concentration was measured according to the manufacturer’s instructions (Mouse Interleukin-6 (IL-6) ELISA Kit; Gentaur, Kampenhout, Belgium). Colorimetric enzymatic tests were performed to measure plasma cholesterol levels (Diasys, Diagnostic System, Holzheim, Germany) following the manufacturer’s instructions.

### Histology and morphological analyses

Five-μm paraffin-embedded kidney sections were stained with Periodic Acid Schiff (PAS), Hemalun, and Luxol Fast Blue to assess morphological alterations. Morphometry of kidney sections was performed as previously reported.^7^ Briefly, the frequency of tubules containing vacuolated cells was evaluated using a semi-quantitative single-blind analysis. To standardize the evaluation procedure, an additional lens engraved with a square grid was inserted into one of the microscope eyepieces. For each paraffin section, 10 square fields (0.084 mm^2^/field) were observed at X400 magnification. Ten randomly selected areas of each cortex kidney section were analyzed using the ImageJ software. Paraffin-embedded liver sections were stained with hematoxylin and eosin and steatosis was graded as described by Ryu and colleagues.^55^

### Western Blot Analysis

Proteins from mitochondrial-enriched renal cortex sample or HK-2 cells were extracted using Cell Lysis Buffer (Cell Signaling, Danvers, MA, USA) with phosphatase and protease inhibitor cocktail (Thermo Fisher Scientific, Waltham, MA, USA) at 4 °C followed by centrifugation at 14,000× g for 15 min at 4°C. Protein concentrations were quantified by Pierce BCA assay kit (Thermo Fisher Scientific, Waltham, MA, USA) and 20 µg of total lysate were separated by SDS-PAGE 12 % and transferred onto nitrocellulose membranes. Membranes were stained with Revert™ 700 Total Protein Stain (Li-Cor Biosciences, Lincoln, NE, USA). Following blocking step in 5 % BSA for 1 h, the membranes were incubated with primary antibodies against acetyl K68-SOD2 (Abcam), SOD2 (Abcam), pan-acetyl lysines (Abcam) or β-actin (Thermo Fisher Scientific, Waltham, MA, USA) overnight at 4°C and then with secondary antibodies (Li-Cor Biosciences, Lincoln, NE, USA) for 1 h at room temperature. Antibodies were diluted in Odyssey Blocking Buffer TBS containing 0.1% Tween 20. Proteins were visualized and quantified using the Odyssey^®^ imaging system (Li-Cor Biosciences, USA).

### Cell culture and treatments

HK-2 cells (Human Kidney 2, ATCC^®^ CRL-2190™, Belgium) were cultivated in DMEM/F12 (Thermo Fisher Scientific, Waltham, MA, USA) supplemented with 10 % fetal bovine serum (FBS, Gibco), 1000 U/L penicillin, 100 µg/mL streptomycin in an atmosphere containing 5% CO2 at 37 °C. Sodium palmitate (Sigma-Aldrich, P9767) was added in 10 % free-FA BSA (Sigma-Aldrich, A8806) solution at 37°C under agitation to reach PA concentration of 7.5 mM stock solution and a PA/BSA molar ratio of 5:2. The concentration was assessed after each complexation using the non-esterified FA assay kit (Fujifilm WAKO, 434-91795 and 436-91995,). HK-2 cells were treated with 300 µM PA or BSA as a vehicle with or without 1 mM Nicotinamide Riboside (NR) (Niagen, ChromaDex, Irvine, CA) for 24h.

### Viability assay

Cell viability was assessed indirectly by staining with crystal violet dye, as described by Journe and colleagues.^56^ Briefly, HK-2 were seeded in 96-well plates at a density of 10.000 cells/well in complete DMEM/F12 medium for 18h. After 24 h treatment, cells were gently rinsed with phosphate-buffered saline (PBS), fixed with 1% glutaraldehyde (Sigma-Aldrich, G7776)/PBS for 15 minutes and stained with 0.1 % crystal violet (Sigma-Aldrich, HT901; weight/vol in double distilled H2O) for 30 minutes. Cells were washed under running tap water for 15 minutes and subsequently lysed with 0.2% Triton X-100 (vol/vol in double distilled H_2_O). The absorbance was measured at 550 nm in a spectrophotometer (SpectraMax ID5, Molecular Devices, USA).

### Metabolic activity

Metabolic activity of cells was assessed by MTT colorimetric assay (Promega, Madison, WI, USA). HK-2 cells were seeded on a 96-well plate at a density of 10.000 cells/well in complete DMEM/F12 medium for 18 h and then treated with BSA/PA during 24 h. Cells were then incubated with MTT (50 μg/well) for 2 h. The supernatant was removed and 100 μl dimethylsulfoxide was subsequently added to each well. After shaking the plate, the absorbance of each well was measured at 570 nm in a spectrophotometer (SpectraMax ID5, Molecular Devices, USA).

### Measurement of lipid droplets content

LD were observed using BODIPY 493/503 lipid probe (Thermo Fisher Scientific, Waltham, MA, USA, D3922). 50.000 cells/well were seeded in 2-well Nunc Lab-Tek Chamber Slide System (Thermo Fisher Scientific, Waltham, MA, USA) for 18 h. After 24 h treatment, cells were washed 3 times with PBS and incubated with 1 mL/well of DMEM/F12 containing 2.5 µM of BODIPY and 3.75 µg/mL of Hoechst 33342 (Miltenyi Biotec, Germany) for 15 min at 37°C. Then, cells were washed 3 times with PBS and immediately examined under a confocal microscope (Leica Microsystems). Quantifications of LD size and number were performed using FIJI v.2.1.0. and the “*MRI Lipid Droplets Tool*” plugin. Quantification was performed on five randomly selected micrographs per condition, with each containing 25 cells. For Oil Red O staining, cells were first rinsed three times with 1X PBS and then fixed with 4% PAF for 30 minutes at room temperature. After fixation, the cells were rinsed three times with PBS, followed by a single rinse with 60% isopropanol. They were then left to dry at room temperature for 10 minutes until the liquid had fully evaporated. The Oil Red O staining solution (Sigma-Aldrich, Saint-Louis, MO, USA) was added, and cells were incubated with the stain for 2 hours at room temperature. After incubation, cells were rinsed four times with PBS. Finally, coverslips with the cultured cells were mounted onto slides using ImmunoHistoMount™ mounting medium (Sigma-Aldrich, Saint-Louis, MO, USA).

### NAD^+^/NADH ratio assessment

NAD^+^ levels were measured using the NAD/NADH-Glo Assay Kit (Promega, Madison, WI, USA) following manufacturer’s instructions. Briefly, cells were seeded on 24-well plate at a density of 50.000 cells/well in complete DMEM/F12 medium for 18 h. Cells were then exposed to treatments for 24h as described before. Cells were washed in PBS and lysed with 220 µL/well of 1 % dodecyltrimethylammonium bromide solution. The content of each well was divided in two to measure NAD+ and NADH separately. To measure NAD+, 50 µL of 0.4 N HCl were added, and cell lysates were heated at 60 °C for 15 minutes. Samples were incubated at room temperature for 10 minutes and 50 µL of Trizma base^®^ were added. To measure NADH, cell lysates were heated at 60 °C for 15 minutes, incubated at room temperature for 10 minutes and 50 µL of HCl/Trizma^®^ solution were added. Finally, 25 µL of cell lysates were loaded in an opaque 96-well plate as well as 25 µL of the NAD+/NADH-Glo™ Detection Reagent containing Ultra-Glo™ Luciferase, Reductase, Reductase Substrate, NAD^+^ Cycling Enzyme and NAD^+^ Cycling Susbtrate. After an incubation of 30 minutes at room temperature, luminescence was recorded using a spectrophotometer (SpectraMax ID5, Molecular Devices, USA).

### Staining of the mitochondrial network

The mitochondrial network was stained through different probes; MitoTracker^TM^ Red CMXRos (Thermo Fisher Scientific, Waltham, MA, USA, M7512), whose accumulation is dependent upon membrane potential and MitoTracker™ Green FM (Thermo Fisher Scientific, Waltham, MA, USA, M7514) whose accumulation is not. In addition, the MitoProbe™ JC-1 (Thermo Fisher Scientific, Waltham, MA, USA, M34152) was used and allowed detection of changes in mitochondrial membrane potential indicated by fluorescence emission shift from green to red.

Briefly, 50.000 cells/well were seeded in 2-well Nunc Lab-Tek Chamber Slide System for 18 h. After 24 h treatment, cells were incubated for 30 min at 37°C with 1 mL/well of DMEM/F12 containing 3.75 µg/mL of Hoechst and 50 nM of MitoTracker^TM^ Red or Green or 200 µM of JC-1. Then, cells were washed 3 times with PBS and immediately examined under a confocal microscope (Leica Microsystems). Mitochondrial network morphology analysis was performed based on MitoTracker™ Green staining. The form factor which is the ratio between end and branched points was calculated with the ImageJ software on 30 randomly selected micrographs containing individual cell per condition. The intensity of fluorescence of MitoTracker^TM^ Red and the JC-1 related fluorescence shift from 590 to 510 nm were calculated using the ImageJ software on five randomly selected micrographs containing 25 cells per condition.

### Assessment of mitochondrial and cytosolic oxidative stress

Mitochondrial superoxide generation was detected using MitoSOX™ Red Mitochondrial Superoxide Indicator (Thermo Fisher Scientific, Waltham, MA, USA, M36008). This dye selectively targets mitochondria, where it is oxidized by superoxide, resulting in red fluorescence. In parallel, lipid peroxidation was assessed by the Image-iT™ Lipid Peroxidation (Thermo Fisher Scientific, Waltham, MA, USA, C10445) which shifts from red to green upon oxidation and thus provides a ratio metric indication of lipid peroxidation.

For both staining, 50.000 cells/well were seeded in 2-well Nunc Lab-Tek Chamber Slide System for 18 h. After 24 h treatment, cells were incubated for 15 min at 37°C with 1 mL/well of DMEM/F12 containing 3.75 µg/mL of Hoechst and 5 µM MitoSOX or 10 nM lipid peroxidation dye. Then, cells were washed 3 times with PBS and immediately examined under a confocal microscope (Leica Microsystems). MitoSOX-related mean fluorescence intensity and the fluorescence shift from 590 to 510 related to lipid peroxidation were obtained through ImageJ from five randomly selected field per condition, with each containing 25 cells.

### DNA, RNA extraction and RT-qPCR

Total DNA extraction from HK-2 cells was performed using the QIAamp DNA Mini Kit (Qiagen, 51304). Following this extraction, an RNase A treatment (Sigma Aldrich, 11119915001) is carried out to degrade any present RNA. Total DNA concentration was measured using NanoDrop 1000 (Thermo Fisher Scientific, Waltham, MA, USA). Total RNA from HK-2 cells was extracted with Trizol (Sigma-Aldrich) and treated with DNAse (Promega). Then, total RNA concentration was measured using NanoDrop 1000 (Thermo Fisher Scientific, Waltham, MA, USA). Reverse transcription was performed with the Transcriptor First Strand cDNA Synthesis Kit (Roche Applied Science, 04897030001) to convert 1 μg of RNA into cDNA. The NCBI Primer BLAST was used to ensure the specificity of the primers for each target. All primer pairs were analyzed for their dissociation curves and melting temperatures. Real-time quantitative PCR was performed to quantify the mRNA levels of *Tfam, Bnip3l, Nd1, Actb* and *18S* as housekeeping gene (**Table 1**). Quantitative PCR amplification was performed using SYBR Green Master Mix (Roche, Belgium) and Prism 7300 Real-Time PCR Detection System (Applied Biosystems, CA, USA). Mean fold changes were calculated by averaging triplicate measurements for each gene. The relative gene expression was calculated using the 2^−ΔΔCT^ method.

**Table 1.**
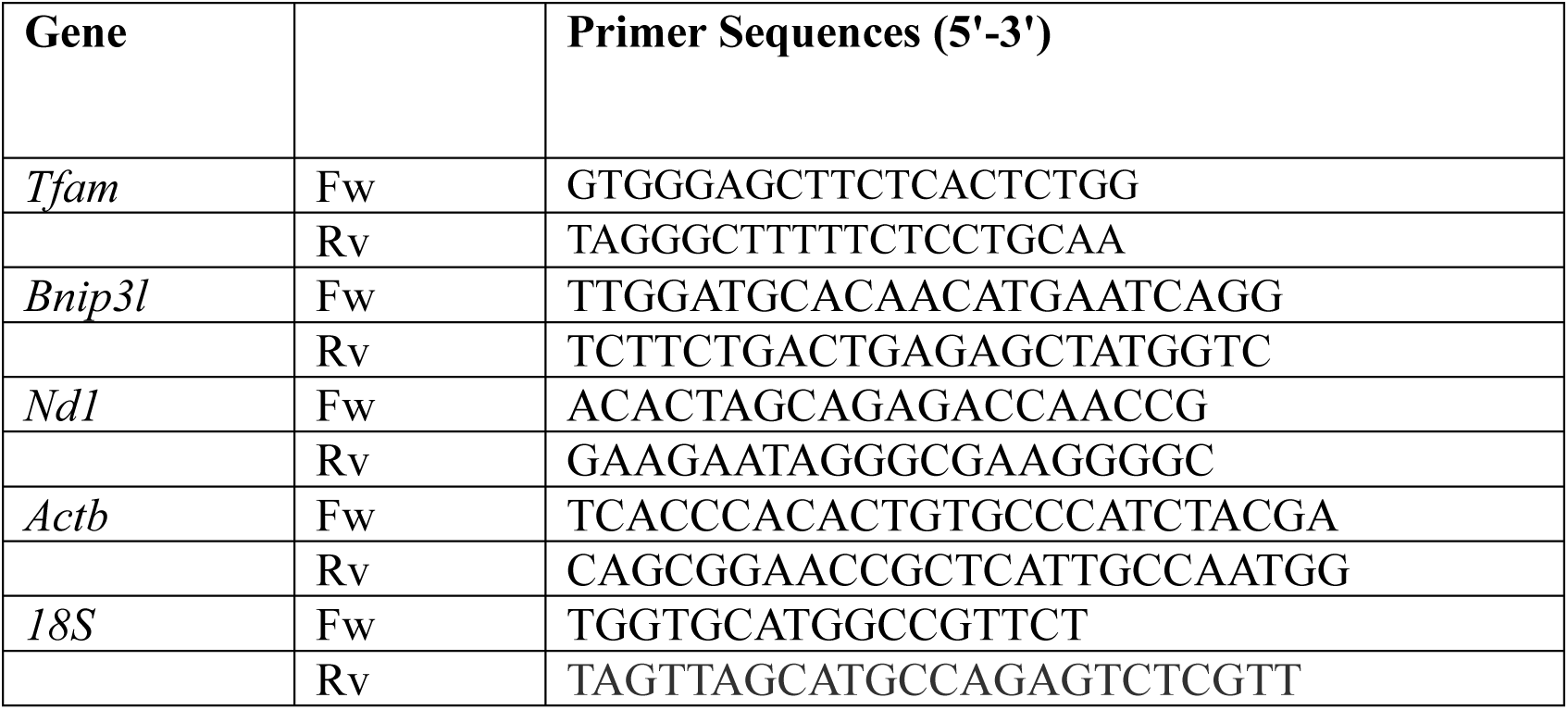

## Statistical analysis

Results are presented as mean values ± SEM. The level for statistical significance was defined as p < 0.05. Analyses were carried out using Prism GraphPad Software version 8. Differences between data groups were evaluated using one-way ANOVA followed by Newman–Keuls *post hoc* tests for multiple comparisons. The level of significance was defined as 0.05 and p values were indicated by symbols in the Figure legends. All experiments were performed at least three times or more.

**Supplementary Figure 1.**
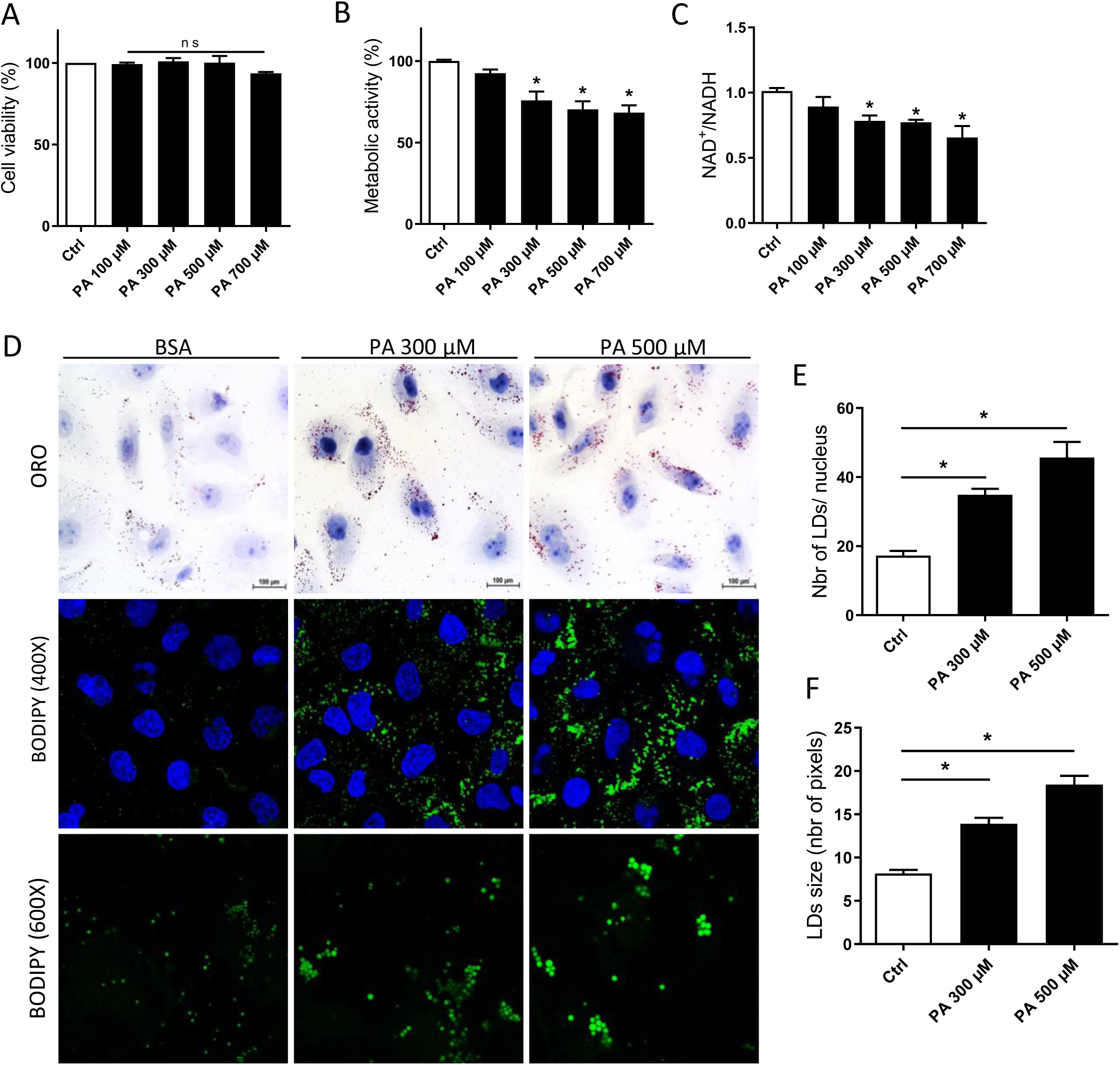
PA induces lipid accumulation and metabolic dysfunction in HK2. **A.** Evaluation of HK-2 cells viability for control, 100, 300, 500 and 700 µM of PA treatment after 24h. **B**. Metabolic activity of HK-2 cells determined with the MTT test for control 100, 300, 500 and 700 µM of PA treatment after 24h. **C.** The NAD+/NADH concentration was measured using the NAD/NADH-Glo Assay Kit in HK2 for control 100, 300, 500 and 700 µM of PA treatment after 24h. **D.** Confocal analysis of BODIPY 493/503 staining (green) to visualize lipid droplets structures in HK2 treated with PA 300 and 500 µM. Nuclei were stained with Hoechst Dye (blue). **E**. Quantifications of number of lipid droplets obtained from five randomly selected field per condition, with each containing 20-30 cells, normalized with the number of nuclei. (400X). n=3. **F**. Quantifications of changes in the average size of lipid droplets in HK2 cells. Each point represents the average size of lipid droplets in a cell. (600X). Data are represented as means ± SEM of three independent biological experiments. Statistical analyses were performed using one-way ANOVA followed by Tukey’s post-hoc test. *p ≤ 0.05

